# Analysis of polyurethane/gelatin complex hydrogel system for protein imprinting

**DOI:** 10.1101/2024.03.03.583225

**Authors:** Alegi Porchkhidze, Madona Endeladze, Nana Gogichaishvili, Otar Mikautadze

## Abstract

Polyurethane served as the carrier in the synthesis of a hydrogel system, incorporating membrane protein as the template, N-Vinylformamide as the monomer, and 1,4-Butanediol diglycidyl ether as the crosslinker, along with gelatin, initiated by ultraviolet radiation. This resulted in the formation of the hydrogel PUNVF-Gelatin. The study investigated the influence of varying monomer concentration, crosslinker concentration, and gelatin concentration on both the adsorption capacity for membrane protein and the imprinting efficiency. Findings revealed that optimal conditions for achieving the maximum adsorption capacity occurred when the monomer mass fraction was 5%, the crosslinker mass fraction was 3%, and the gelatin mass fraction was 0.6%.

## 1. Introduction

Hydrogels, with their versatile properties, play a pivotal role in scientific applications, notably in molecular imprinting. Their unique advantage lies in their ability to imprint large biomolecules like proteins while minimizing interference with native structures.[1-5] In the realm of molecular imprinting, hydrogels delicately balance molecular recognition and preservation of the native protein structure, making them crucial for success.[6, 7] This synergy between hydrogels and molecular imprinting not only advances technologies like drug delivery and biomimetic sensors but also promises innovative solutions in personalized medicine, incorporating statistical and machine learning methods.[8-10] The marriage of hydrogel versatility and molecular precision opens new frontiers in tailored recognition materials, enriching our understanding of biomolecular interactions and impacting various scientific and biomedical disciplines.[11-13]

N-Vinylformamide (NVF) hydrogels, formed through free radical polymerization, exhibit excellent water absorbency and responsiveness to stimuli like temperature and pH changes.[14-16] These versatile hydrogels hold promise for controlled drug delivery, wound healing, and biomedical applications, with the potential for statistical and machine learning optimization.[17, 18] Notably biocompatible, NVF hydrogels are actively explored for use in tissue engineering, regenerative medicine, and other fields due to their mimicry of the natural extracellular matrix.[19-21] With their unique properties, NVF hydrogels present exciting opportunities for innovation in various scientific and biomedical applications.

Gelatin-based hydrogels, derived from collagen, are versatile biomaterials with excellent biocompatibility.[22-24] These hydrogels mimic the extracellular matrix, promoting cell adhesion and tissue regeneration.[25, 26] Tunable mechanical properties and controlled degradation make them ideal for applications in tissue engineering, incorporating statistical and machine learning models for optimization.[27, 28] Gelatin hydrogels are also utilized for controlled drug delivery due to their encapsulation capabilities and responsiveness to environmental stimuli.[29, 30] With these unique qualities, gelatin-based hydrogels continue to drive innovation in areas like wound healing, drug delivery, and tissue engineering.[31, 32]

The evolving field of hydrogel-based imprinting techniques also extends to advanced methodologies such as surface imprinting, where statistical and machine learning algorithms can enhance design precision.[33-38] Techniques involving the imprinting of molecules on the surface of inorganic or organic carriers have gained prominence due to the resulting materials exhibiting well-defined morphology, thermal stability, and mechanical strength.[39, 40] This growing area of research opens new possibilities for designing hydrogel-based materials with enhanced imprinting efficiency and stability through the integration of statistical and machine learning methods.

In the context of this study, our approach involves utilizing polypropylene (PU) as a carrier, further expanding the repertoire of materials compatible with hydrogel-based imprinting. The synergy of PU, membrane protein as the template, N-Vinylformamide (NVF) as the monomer, and 1,4-Butanediol diglycidyl ether (BDDE) as the cross-linker, initiated by ultraviolet radiation in the presence of gelatin, presents a unique avenue for creating PU-grafted membrane protein-imprinted NVF/gelatin hydrogels. This study not only delves into the intricacies of monomer concentration, cross-linker concentration, and sodium alginate concentration but also seeks to unravel their collective impact on protein adsorption and imprinting efficiency, employing statistical and machine learning analyses. Through a comprehensive exploration of these parameters, we aim to contribute to the evolving landscape of hydrogel-based imprinting techniques, providing insights that may pave the way for innovative applications in areas such as biotechnology, pharmaceuticals, and biomaterials.

## 2. Experiments

### 2.1 Materials

Polyurethan (PU), membrane protein, N-Vinylformamide (NVF), 1,4-Butanediol diglycidyl ether (BDDE), gelatin Acetic acid and Sodium dodecyl sulfate (SDS) were purchased from Sigma.

### 2.2 Procedures

The preparation of PU-NVF-Gelatin hydrogels is described as follows: Take 50 mL of membrane protein solution with a concentration of 2.5 mg/mL. Add different mass fractions (5%, 10%, 15%, 20%, 30%) of NVF, cross-linker BDDE at mass fractions of 0.5%, 1%, 2%, 3%, 5%, 10% a photo initiator, and various mass fractions (0, 0.25%, 0.5%, 1%, 1.5%, 2.0%) of gelatin, stirring until fully dissolved. Immerse pre-weighed PU in the 20 mL solution for 12 hours. After soaking, place the PU on a glass plate, purge with nitrogen, seal, and irradiate with a 100 W UV lamp for 30 minutes. Subsequently, immerse it in a 5.0% calcium chloride solution for 3 hour for cross-linking, resulting in PU-NVF-Gelatin hydrogels. Prepare a 1% SDS in 10% acetic acid solution as the eluent. Place the protein-containing PU grafted hydrogel into 30 mL of the eluent. Shake at a constant temperature (100-200 rpm) in an orbital shaker for 24 hours to elute the template protein, obtaining control hydrogels.

### 2.3 Calculation of Grafting Rate

The grafting rate is calculated based on the increased mass of PU after grafting. The prepared grafted PU is vacuum-dried at 50-70°C until a constant weight is achieved. The grafting rate GR is calculated using equation (1).

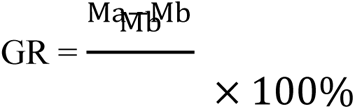

Where Ma and Mb are the masses of PU before grafting and after grafting, respectively.

### 2.4 Calculation of Adsorption Capacity and Imprinting Efficiency

The water on the surface of the PU-NVF-Gelatin and control is dried with filter paper, and approximately 1 g is weighed and placed separately in 10 mL of membrane protein absorption solution with a mass concentration of 1.5 mg/mL for soaking for 24 hours. The UV absorbance of the BSA solution at 278 nm is measured using a UV-visible spectrophotometer. The mass concentration of before and after membrane protein adsorption is obtained according to the standard curve, and the adsorption quantity QA is calculated using equation (2).

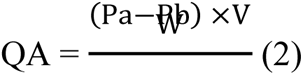

Where Pa is the initial mass concentration of the membrane protein solution; Pb is the mass concentration of the membrane protein solution at a certain time, mg/mL; V is the volume of the membrane protein solution, L; W is the mass of the grafted PU, g; The imprinting efficiency E is calculated according to equation (3):

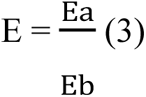

Where Ea is the equilibrium adsorption quantity of PU-NVF-Gelatin and Eb is the adsorption quantity of control.

## 3. Results and Discussion

### 3.1 Impact of Monomer Mass Fraction on Grafting Yield

Figure 1 illustrates the curve of grafting yield with the variation of monomer mass fraction. In this study, the cross-linking agent mass fraction was fixed at 1%, while monomers with different mass fractions were prepared to investigate their influence on grafting yield. As shown in Figure 1, with the increase in monomer mass fraction, the grafting yield rapidly increases until the monomer mass fraction reaches 25%, at which point the grafting yield stabilizes. In the early stages of the reaction, the grafting yield increases rapidly with the rise in monomer mass fraction because a higher monomer mass fraction leads to the involvement of more monomers in the grafting reaction. However, when the monomer mass fraction reaches 25%, the self-polymerization probability of the monomers increases, causing a rapid increase in system viscosity, which hinders the diffusion of free radicals, leading to a decrease in grafting yield.

**Figure 1.**
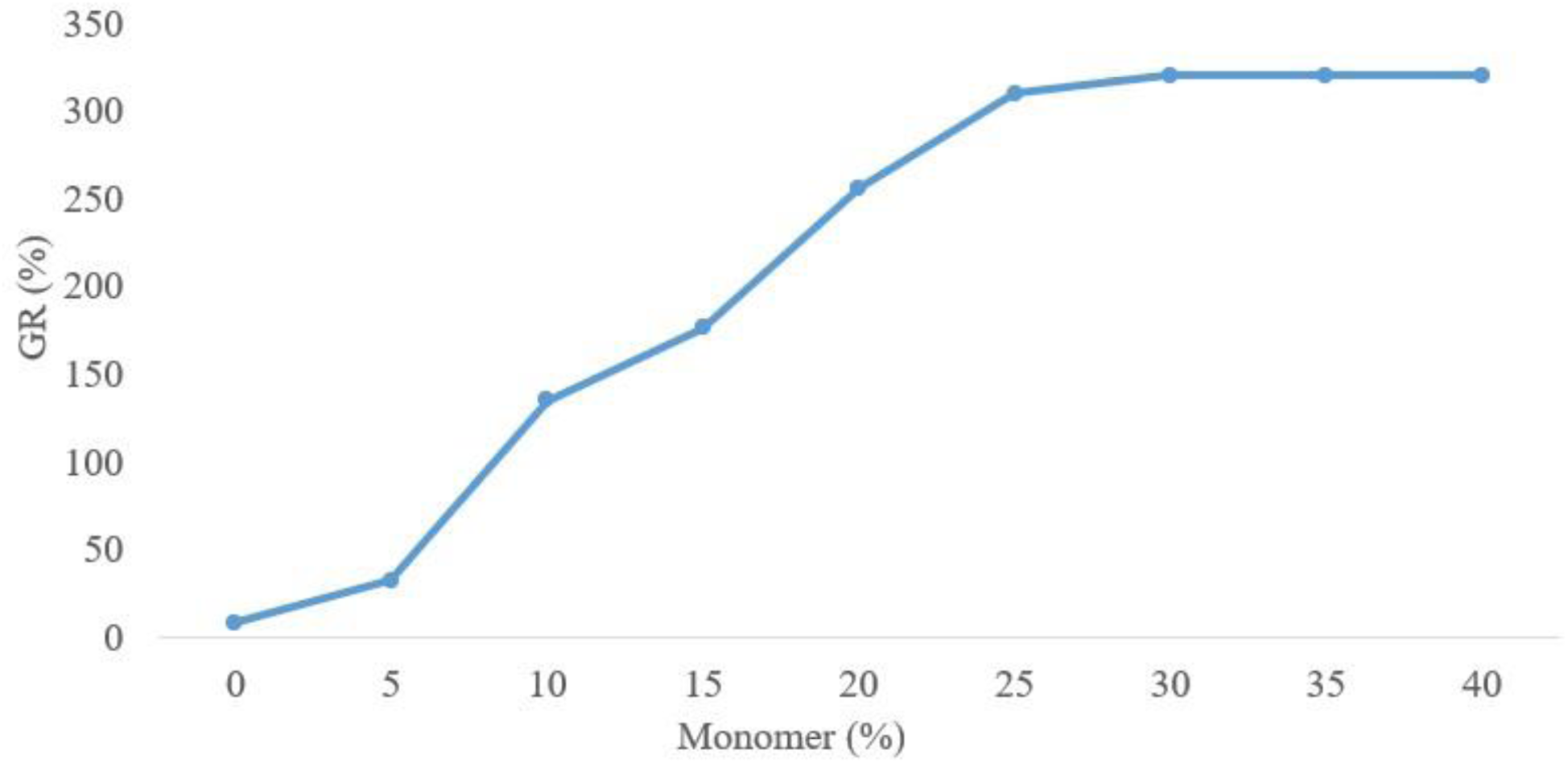

### 3.2 Influence of Monomer Mass Fraction on Membrane Protein Adsorption and Imprinting Efficiency

Under fixed experimental conditions (cross-linking agent mass fraction of 1%, gelatin mass fraction of 0.5%), Figure 2 shows the adsorption amounts of PU-NVF-Gelatin and control on membrane proteins at different monomer mass fractions. When the monomer mass fraction is 5%, PU-NVF-Gelatin exhibits the maximum protein adsorption; however, the adsorption rapidly decreases when the monomer mass fraction exceeds 5%. In contrast, control gradually reaches equilibrium in adsorption after a monomer mass fraction greater than 5%. At lower monomer mass fractions, the prepared material facilitates protein diffusion, and increasing the monomer mass fraction can generate more imprinting sites and pores. As the monomer mass fraction increases, the prepared material becomes denser, increasing spatial resistance for membrane proteins, making protein desorption more difficult and resulting in a decrease in adsorption. From Figure 2, it can be observed that PU-NVF-Gelatin has a higher adsorption capacity for membrane proteins compared to control. Figure 3 demonstrates the imprinting efficiency of PU-NVF-Gelatin on membrane proteins at different monomer mass fractions. It is evident that the imprinting efficiency reaches its maximum value at a monomer mass fraction of 10%, which is 4.2.

**Figure 2.**
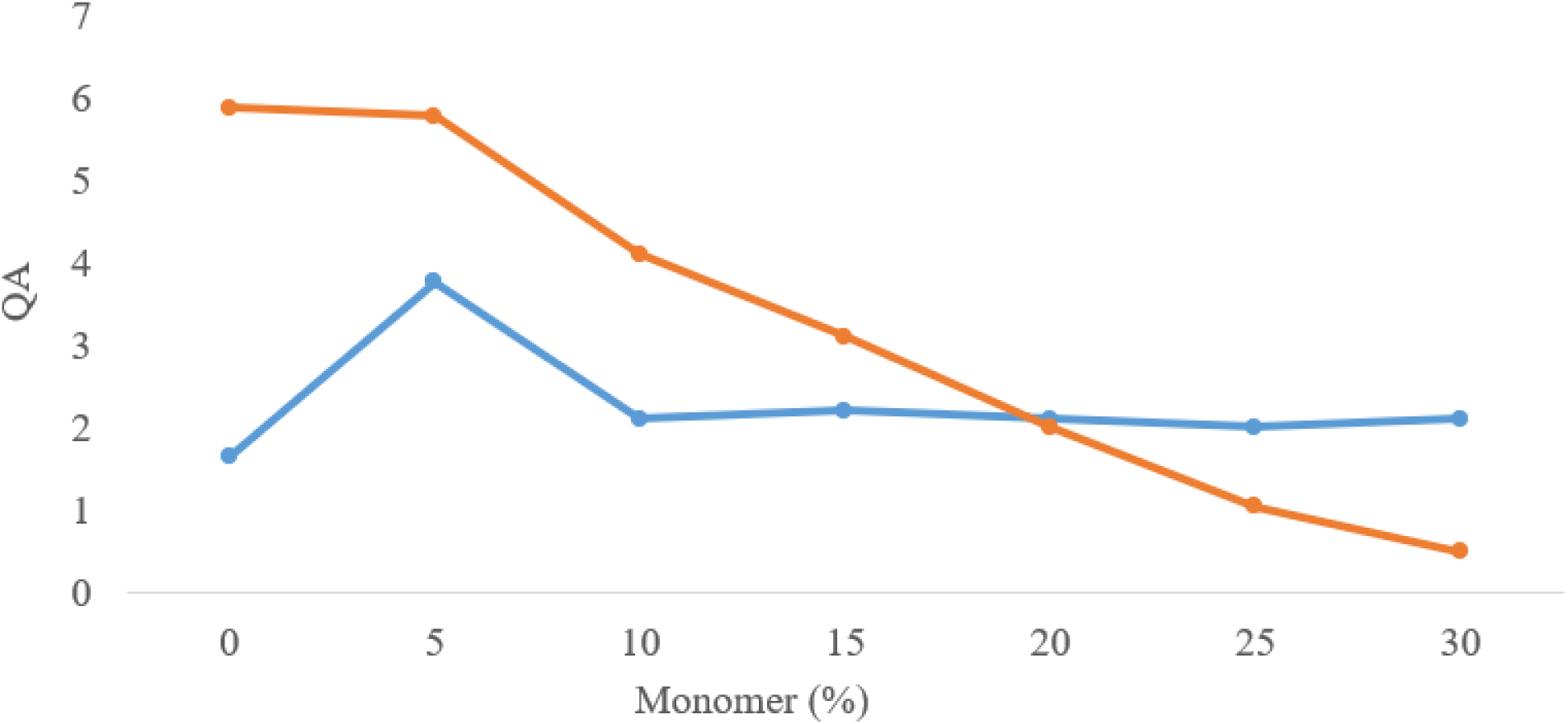

**Figure 3.**
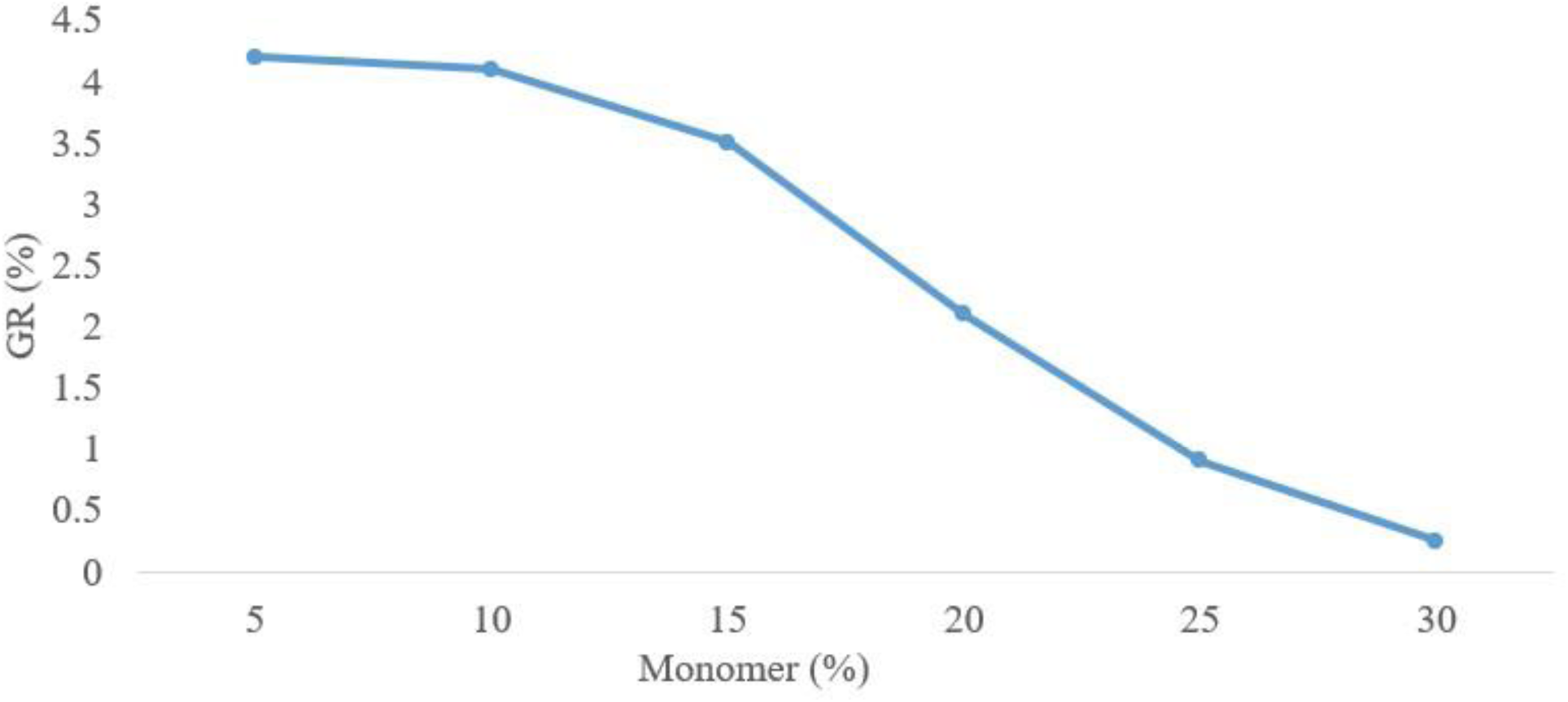

### 3.3 Impact of Cross-Linking Agent Mass Fraction on membrane Protein Adsorption and Imprinting Efficiency

Under fixed experimental conditions (monomer mass fraction 10%, gelatin mass fraction 0.5%), Figure 4 reveals that as the cross-linking agent mass fraction increases from 1% to 5%, the adsorption of PU-NVF-Gelatin on membrane proteins gradually rises. However, after the cross-linking agent mass fraction exceeds 3%, the adsorption rapidly decreases. This is because in molecular imprinting materials, the polymer matrix is typically a highly cross-linked (cross-linking degree > 90%) rigid material to maintain the shape of imprinting sites. Highly cross-linked rigid materials cannot imprint large and flexible membrane proteins due to their small internal pores. At lower cross-linking agent mass fractions, the formed gel is too soft to create effective binding sites. With an increase in the cross-linking agent mass fraction, the gel strength increases, forming stable imprinting sites, and the adsorption amount increases. However, when the cross-linking agent mass fraction further increases, the spatial mesh of the gel decreases, hindering the mass transfer of membrane proteins into the gel, resulting in a decrease in adsorption. From Figure 5, it can be observed that with the increase in the cross-linking agent mass fraction, the imprinting efficiency rapidly increases, reaching its maximum value of 3.75 at a cross-linking agent mass fraction of 2%. Subsequently, the imprinting efficiency gradually decreases. This is because lower cross-linking degrees lead to excessively soft gels that cannot form effective imprints, and increasing the cross-linking agent mass fraction can enhance the number of effective imprinting sites, thereby improving imprinting efficiency. However, excessively high cross-linking degrees also hinder the entry and exit of membrane proteins from the gel, reducing imprinting efficiency.

**Figure 4.**
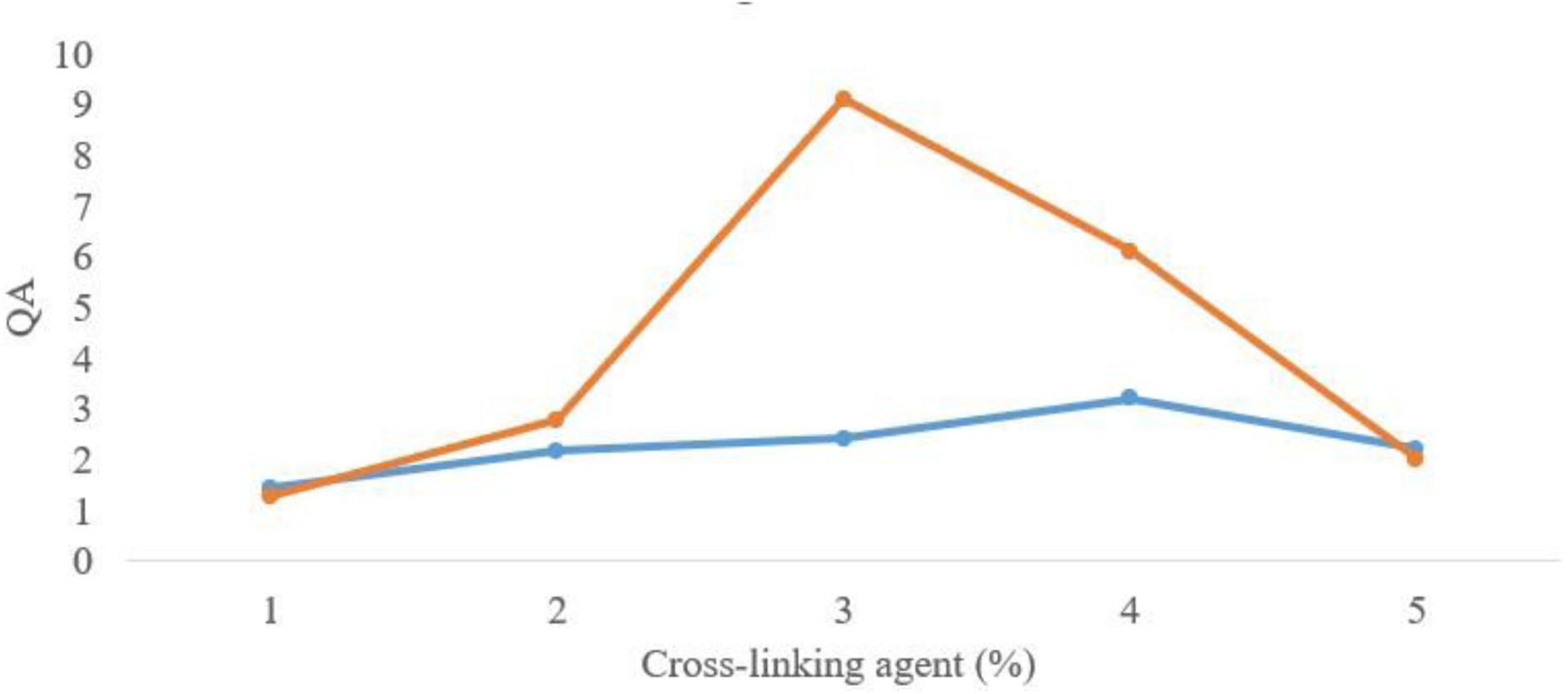

**Figure 5.**
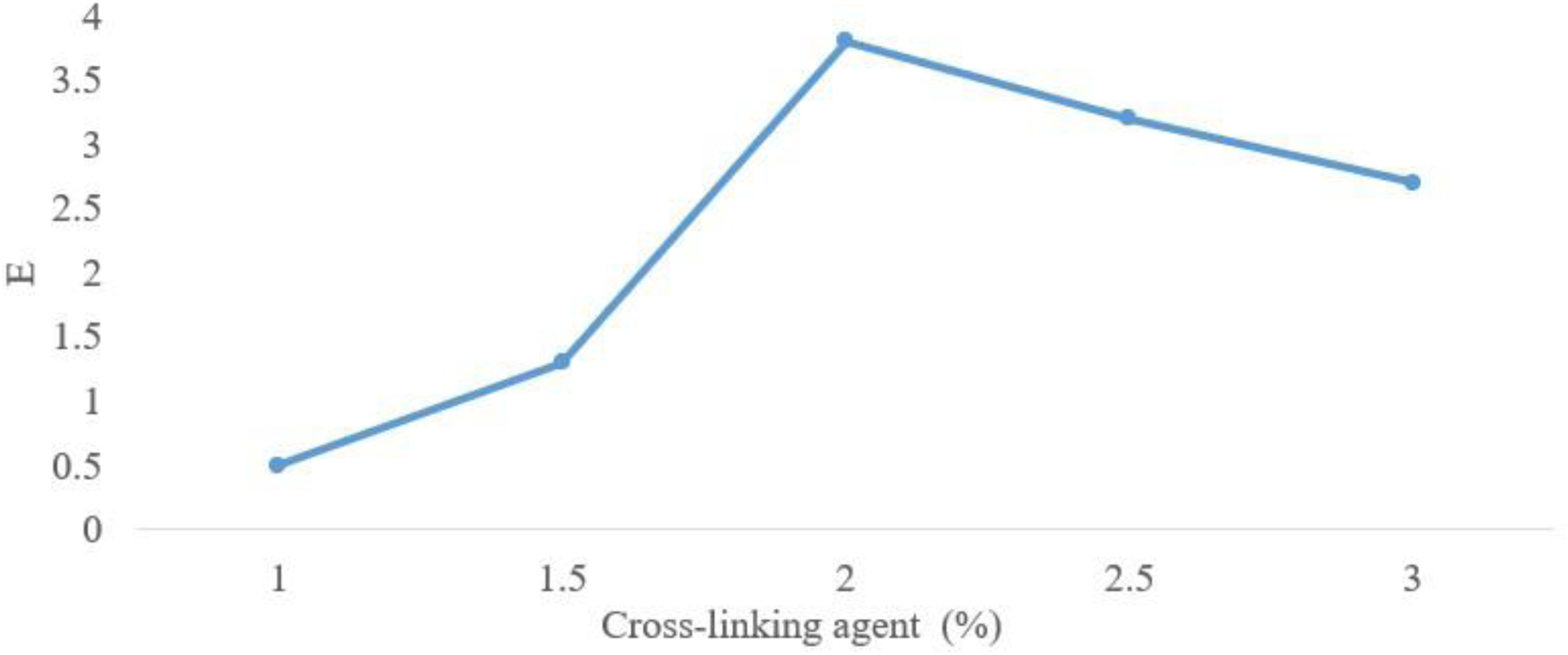

### 3.4 Impact of Gelatin Mass Fraction on membrane Protein Adsorption

Under fixed experimental conditions, specifically a monomer mass fraction of 15% and a cross-linking agent mass fraction of 1%, Figure 6 reveals that PU-NVF-Gelatin has the maximum adsorption capacity when the sodium alginate mass fraction is 0.5%. When the Gelatin mass fraction is below 0.6%, the adsorption of membrane proteins rapidly increases with an increase in Gelatin mass fraction. However, when the Gelatin mass fraction exceeds 0.6%, the imprint adsorption capacity gradually decreases. This is because the addition of Gelatin forms a cross-linked network structure, increasing the number of imprinting sites. At lower Gelatin mass fractions, the carboxylate ions, ionized in the aqueous solution, can interact with amino groups in membrane proteins, facilitating the re-binding of large molecules of membrane proteins to complementary sites in the polymer, resulting in an increase in adsorption. However, when the Gelatin mass fraction reaches 0.6% and beyond, an increase in Gelatin mass fraction leads to a reduction in gel pore size, impeding the mass transfer of membrane proteins into the gel and causing a decrease in adsorption capacity. On the other hand, with an increase in Gelatin mass fraction, the carboxylate ion content in the gel increases. When it reaches a certain level, the electrostatic repulsion between the carboxylate ions and the carboxyl groups in membrane proteins becomes greater than their binding with amino groups. Consequently, this leads to a decrease in adsorption capacity.

**Figure 6.**
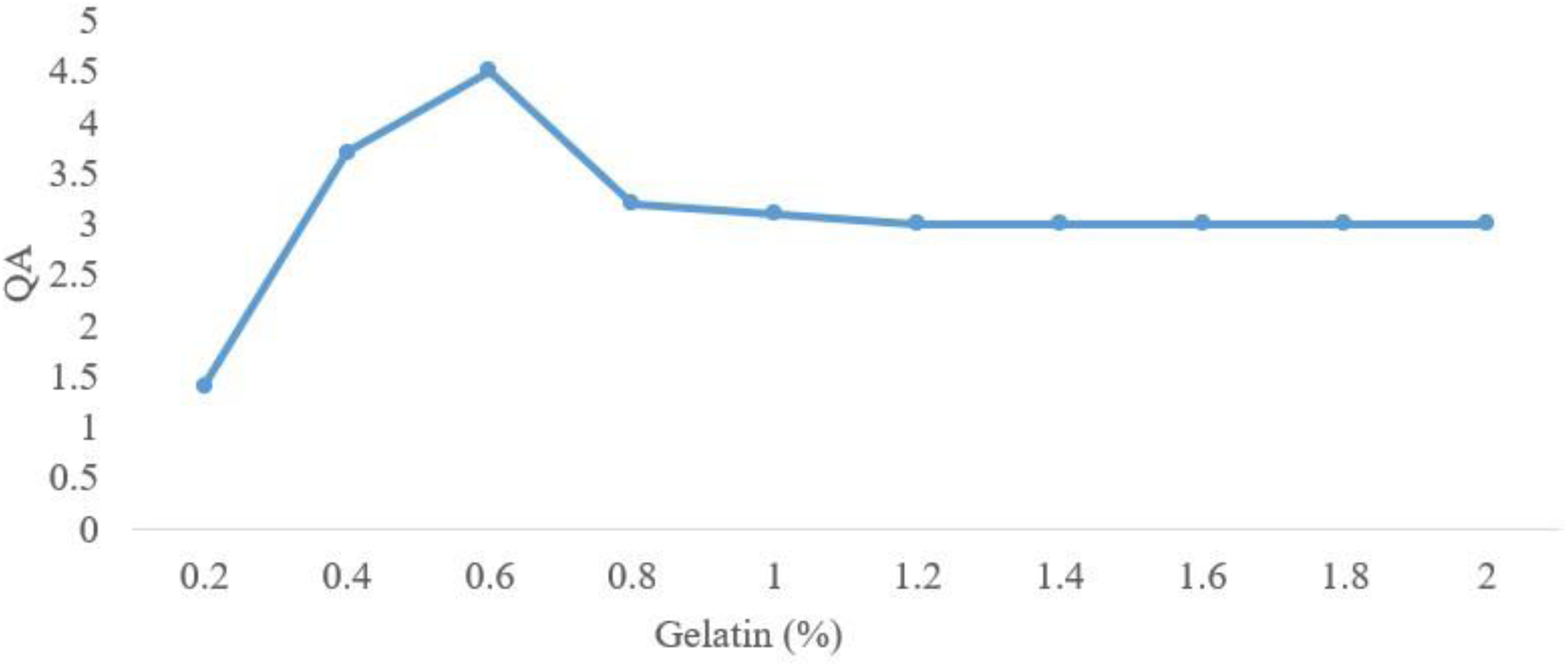

## 4. Conclusions

From this study, we concluded that PU as the carrier, membrane protein as the template, NVF as the monomer, and BDDE as the cross-linking agent, in the presence of gelatin and initiated by ultraviolet radiation, PU can be grafted with, membrane protein-imprinted NVF/gelatin hydrogel (PU-NVF-Gelatin). When the monomer mass fraction is 5%, the cross-linking agent mass fraction is 3%, and the gelatin mass fraction is 0.6%, the imprinted hydrogel grafted on PU exhibits the maximum adsorption capacity for membrane protein. PU-NVF-Gelatin demonstrates higher adsorption capacity for the template protein compared to control. The imprinting efficiency reaches its maximum, 3.75, when the cross-linking agent mass fraction is 2%.

